# Sleep’s impact on emotional memory: A meta-analysis of whole-night, nap, and REM sleep effects

**DOI:** 10.1101/732651

**Authors:** Sarah K. Schäfer, Benedikt E. Wirth, Marlene Staginnus, Nicolas Becker, Tanja Michael, M. Roxanne Sopp

## Abstract

Numerous studies have shown that sleep enhances the consolidation of episodic memory. However, it remains unclear whether this consolidation benefit is moderated by the emotional valence of the learned material. To clarify whether sleep selectively enhances the consolidation of emotional material, we conducted a meta-analysis including *N* = 1,059 observations. Overall, our results do not support this hypothesis. When only studies with a sleep-group wake-group comparison were included in the analysis (*k* = 22), the retention advantage for emotional over neutral material was not significantly different between sleep and wake groups. When studies initially lacking a wake-control group were included in the analysis after statistical estimation of wake-group parameters, the retention advantage for emotional material was significantly larger in wake-groups than in sleep-groups (*k* = 34). Interestingly, however, an additional analysis of *k* = 8 studies investigating the selective effects of rapid-eye-movement sleep and slow-wave sleep on emotional memory consolidation provided evidence for a selective enhancement of emotional over neutral memory consolidation after rapid-eye-movement sleep compared to slow-wave sleep. These results suggest that sleep does not generally enhance emotional memory consolidation over neutral memory consolidation. However, specific sleep stages might preferentially enhance consolidation of emotional and neutral material, respectively.

## 1. Introduction

Research of the past two decades has established that sleep plays a crucial role in memory consolidation [1]. Models of systems consolidation [2] assume that newly encoded memory traces are reactivated during sleep, which enables the redistribution of memory representations from intermediate-term storage in the hippocampus to neocortical long-term stores. This process is assumed to prevent the decay of memory representations that would occur if storage remained to be supported by hippocampal circuits [3]. In line with this framework, numerous studies show that memory performance is significantly enhanced when encoding is followed by sleep rather than wakefulness (see e.g., [4–6]). These effects are evident after nap [7] and nighttime sleep [8] and have been demonstrated for declarative and procedural memory performance [9]. However, effects are especially pronounced in tests of episodic memory, which is consistent with the functional role of the hippocampus in this memory domain [10,11].

Beyond these behavioral accounts, research has provided insights into the neurophysiological mechanisms supporting memory consolidation during sleep. Sleep is characterized by the cyclic alteration of rapid-eye-movement sleep (REMS) and non-rapid-eye-movement sleep (NREMS). NREMS, in turn, consists of three stages that progressively increase in depth (N1, N2, and slow-wave sleep; SWS). During SWS, neocortical slow oscillations are assumed to drive the reactivation of hippocampal memory traces as reflected in the occurrence of hippocampal sharp wave ripples [2]. These sharp wave ripples are temporally synchronized with thalamocortical sleep spindles, which may mediate the process of memory redistribution [12,13]. Consistent with this model, studies demonstrate that consolidation benefits for episodic memory are stronger after early night sleep dominated by SWS than after late night sleep dominated by REMS [14]. Studies further suggest that reactivations can be triggered by presenting olfactory [15] or auditory [16] memory cues during SWS but not during other sleep stages [15].

Beyond its general role in memory consolidation, sleep has been associated with the consolidation of emotional memories. Research demonstrates that emotional memories are preferentially retained across time [17]. Notably, this effect appears to be enhanced by sleep [18]. Selective effects of sleep on emotional memory have mostly been studied in the context of episodic memory [19]. In these studies, participants are presented with emotional and neutral memoranda followed by a phase of sleep or wakefulness. Subsequently, participants are subjected to a memory test that requires recall or recognition of the memoranda. By contrasting the relative retrieval advantage for emotional (vs. neutral) memoranda between sleep and wake conditions, studies investigate the selective benefits of sleep for emotional memory consolidation. Such studies demonstrated selective benefits of sleep for (mostly negative) emotional memories [20–23]. These benefits were found after nap [24,25] and nocturnal sleep [22,26]. Moreover, studies investigating consolidation benefits across early night SWS and late night REMS indicate that emotional memory performance is particularly high after late night REMS [20,21]. Correspondingly, emotional memory performance is more strongly impaired by REMS deprivation than by SWS deprivation [23].

Based on these findings, theoretical accounts propose that selective reactivation and consolidation of emotional memories may specifically occur during REMS [27]. During REMS, emotional memories are assumed to be reactivated as indicated by enhanced activation of the limbic system [28]. In terms of oscillatory activity, reactivations could be reflected by the propagation of theta oscillations [29]. Theta oscillations partially arise from synchronized activation in the neocortex, hippocampus, and the amygdala [30]. Critically, REMS theta coherence between these structures has been found to predict fear memory consolidation in rats [31]. Moreover, experimental suppression of REMS theta activity in mice is associated with the disruption of fear memory consolidation across sleep [32]. Correspondingly, human studies have linked frontal REMS theta power with subsequent emotional memory performance [24,26].

Despite these converging findings, several studies failed to establish associations between REMS physiology and emotional memory performance [33–36], leaving ambiguities regarding the role of REMS in emotional memory consolidation. Critically, a considerable number of studies did not even replicate selective retention benefits of emotional memories across sleep [37–39]. For instance, Baran et al. [40] and Cairney et al. [41] found emotional and neutral memories to be equally consolidated across sleep. These inconsistencies may result from the variety of study designs used to investigate sleep-related effects (e.g., sleep deprivation, nap sleep, nocturnal sleep, and split-night studies), which vary widely in their degree of experimental control (e.g., lab-monitored sleep, home-monitored sleep, and no sleep monitoring). Moreover, studies employ different variants of recall and recognition tasks (e.g., recollection- and familiarity-based recognition) that use different stimulus materials (e.g., manipulating valence and/or arousal), presentation modes (e.g., number of presented stimuli, duration of stimulus presentation), and task instructions (e.g., incidental vs. explicit encoding). Finally, studies differ in their indices of memory performance (e.g., hit rates, *P_r_*, *d’* or relative retention scores) and data analysis strategies (e.g., analyses including all responses or only recollection/familiarity-based responses). Moreover, the majority of studies suffers from limited statistical power due to small sample sizes [42].

Taken together these methodological issues have raised doubts regarding the reliability of selective emotional memory consolidation during sleep. Ultimately, this ambiguity can only be resolved by estimating the population effect size based on available studies. A recent meta-analysis that attempted to estimate this effect size did not find an overall difference between emotional and neutral memory retention after sleep and wakefulness [43]. However, this null-effect may be related to methodological and statistical shortcomings of the analysis. First, the meta-analysis did not estimate the effect of sleep versus wakefulness on the difference between emotional and neutral memory performance. Instead, the authors conducted separate meta-analyses for wake and sleep groups and compared the magnitude of estimated effect sizes between these conditions. This approach treats emotional and neutral memory performance as independent measures and ignores their substantial within-subject correlation, thus potentially yielding biased effect size estimates. Secondly, the authors aggregated statistically non-equivalent outcome measures (e.g., post-sleep memory scores and pre-post-change scores) from various study designs (e.g., recall tests and recognition tests). Due to these methodological limitations, we investigated the effect of sleep on emotional memory performance adopting a different meta-analytical approach. First, in line with the majority of studies published in this field, we limited our analyses to recognition memory. As memory models [10] assume that recognition tests engage different memory processes than recall tests, we deemed this approach more appropriate than aggregating effect size measures across recognition and recall tests. Second, given that outcome measures differ considerably among studies, we converted them into a common measure (i.e., post-sleep/wake difference between emotional and neutral memory performance). Third, we obtained the within-subject correlations between neutral and emotional memory performance from as many authors as possible and estimated these correlations for the remaining studies. Fourth, to maximize the pool of available studies, we developed a statistical approach that estimates wake group effects for studies without such a group (35% of included studies). Overall, our strategy was tailored to obtain a reliable estimate of the population effect while accouting for procedural differences between studies in subsequent moderator analyses. Moreover, based on the proposed role of REMS in emotional memory consolidation, we conducted an additional meta-analysis investigating the effects of post-learning REMS-rich sleep (vs. SWS-rich sleep) on the difference between emotional and neutral memory performance.

## 2. Methods

### 2.1 Literature search

This meta-analysis was prepared in accordance with the Preferred Reporting Items for Systematic Reviews and Meta-Analyses (PRISMA) guidelines [44] and was pre-registered in the PROSPERO database (ID: CRD42018083899). Relevant search terms were defined to cover the most common terms in the literature on sleep and emotional memory. Using these terms, a literature search based on title, abstract, and keywords was conducted in three databases: EBSCOhost (PsycINFO and PsycARTICLES), Web of Science, and PubMed. In addition, Google Scholar was used to identify publications that cited records that were identified as relevant using the aforementioned databases. No date-of-publication criterion was defined, and all databases covered publication years dating back to at least 1945. Search terms, search engines, as well as the original hits per search engine are displayed in Figure 1. The literature search started on January 8^th^ 2018 and corresponding alerts were followed up until May 1^st^ 2019. The project was further advertised on relevant social-media platforms to increase the likelihood of including unpublished data.

**Figure 1.**
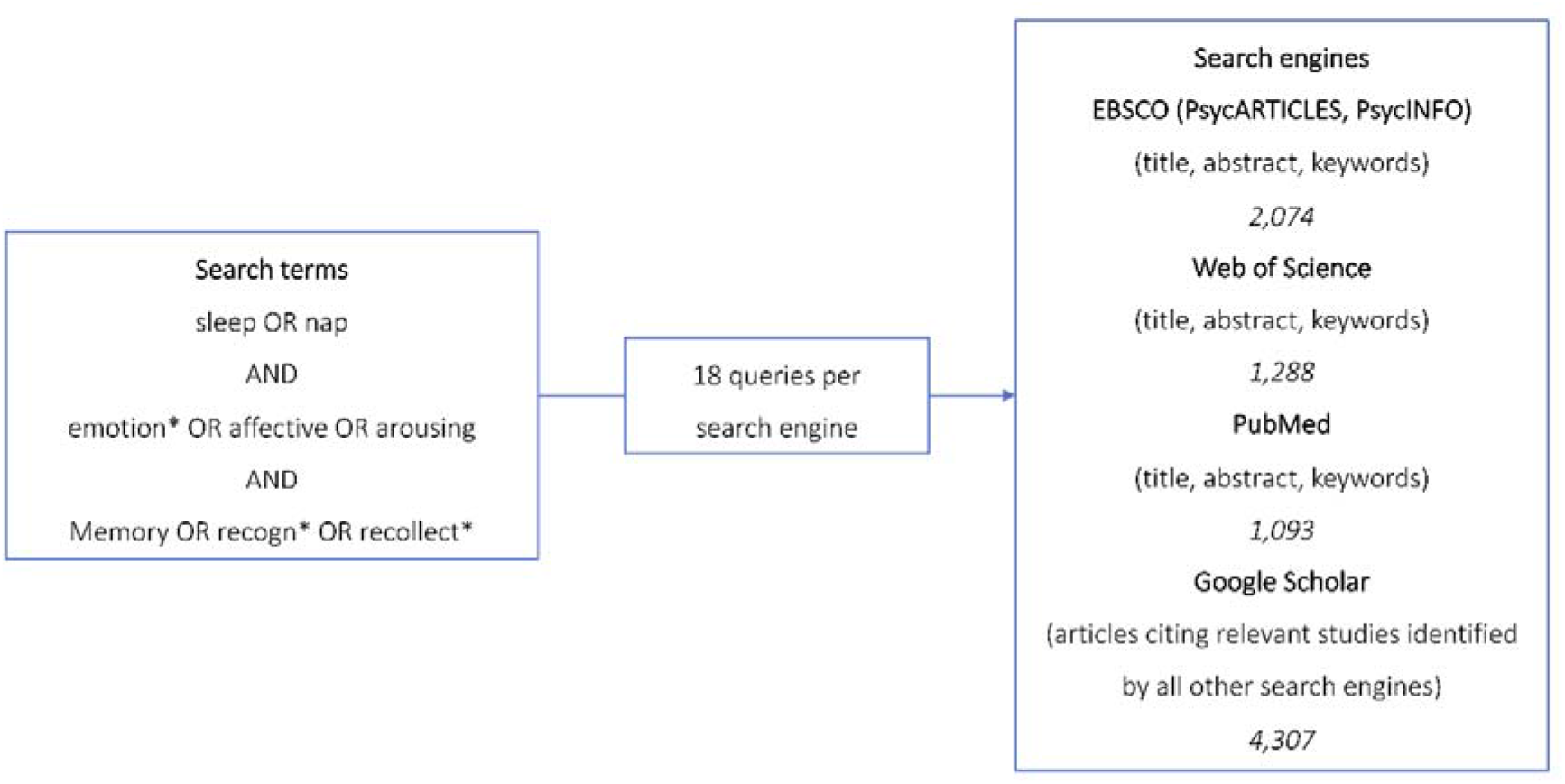
Search terms, search engines, and hits per database.

### 2.2 Inclusion criteria

Samples that met the following criteria were included: 1) The (pseudo-)randomized controlled study reported on a sample that encoded a set of emotional (positive and/or negative) and neutral discrete stimuli (i.e., pictures, words). 2) One group of participants subsequently underwent a post-encoding sleep opportunity. 3) Following sleep and within 48 hours of encoding, participants performed a recognition memory test. 4) Participants were healthy adults aged between 18 and 60 years, a previously used cut-off for old age in sleep research [9]. 5) The study was published in English.

Some studies identified by the literature search did not include a wake control group. To increase the number of eligible samples we decided to include these studies and to estimate missing wake-control groups based on a sub-meta-analysis on available wake-control groups separately for nap and whole night designs. All results are presented both including and excluding these samples.

### 2.3 Exclusion criteria

Samples were excluded if 1) they exclusively investigated the effect of sleep in the context of additional memory tasks (e.g., directed forgetting task) or sleep manipulations (e.g., pharmacological agents), 2) data necessary for effect size calculation was not available by June 9^th^ 2019, 3) for wake samples: participants were sleep-deprived without a recovery night prior to memory testing^1^. Split-night or selective (REMS/SWS) sleep deprivation studies were analyzed in an additional meta-analysis since they do not compare sleep with wakefulness (see 3.8).

### 2.4 Study selection

The study selection process is illustrated in the PRISMA flowchart in Figure 2. Three junior researchers (MS and two graduate students) screened titles, abstracts, and keywords for eligibility. On a trial run of 200 records, a high average inter-rater agreement (94%) was achieved for inclusion/exclusion decisions. After abstract screening, the full texts of 105 records were independently assessed for inclusion eligibility by pairs of junior and senior researchers (BEW, MRS, and SKS), resulting in 51 potentially eligible studies. Of these, 40 studies provided sufficient information to be included, comprising 34 samples for the metaanalysis on nap/whole night designs and 8 samples for the meta-analysis on split-night/REM/SWS deprivation designs.

**Figure 2.**
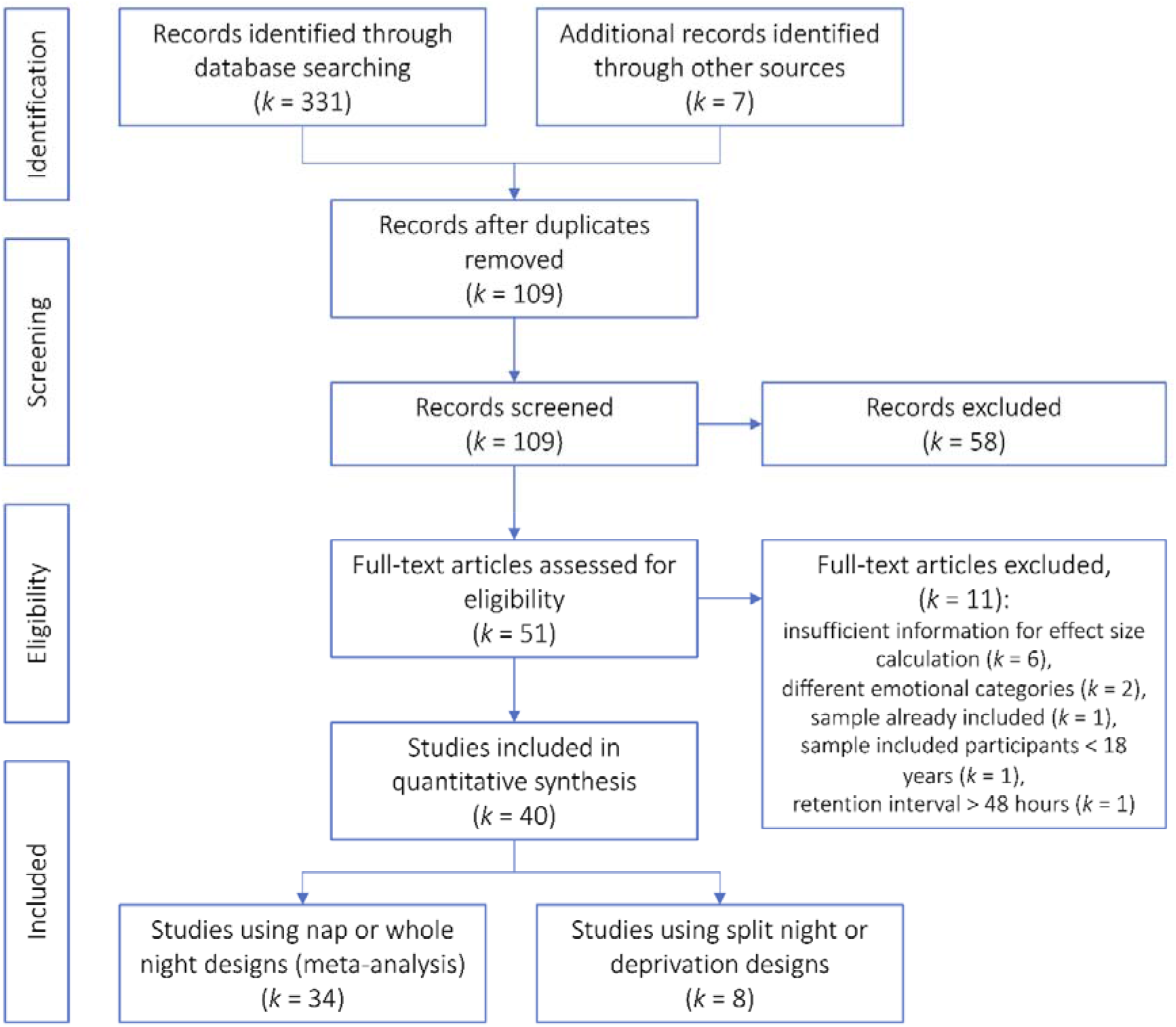
Flowchart of the study selection process according to PRISMA guidelines. One study contributed to both the meta-analysis on nap/whole night designs and split-night/REM/SWS deprivation designs.

### 2.5 Data extraction

Using a standardized Excel form, data for each study was extracted by two independent coders, one junior and one senior researcher, to reduce the likelihood of errors [46]. If available, mean values of corrected memory accuracy measures (i.e., P_r_ and *d’*) were coded to account for response biases. If both P_r_ and *d’* were reported, P_r_ was noted since it is not based on any specific assumptions about underlying memory processes. Hit rates were only included in one case [47] where other measures were not available. The interrater-agreement concerning the extracted data was high: 100% for *n*, 99.4% for *Ms* and 99.72% for *SD*s/*SEM*s. If data necessary for effect size calculation was not reported, the corresponding authors were contacted. If we did not receive an answer to our e-mail or the authors were not able to provide the relevant information, the online tool *WebPlotDigitizer* [48] was used to extract data from figures. All disagreements between coders were resolved through discussion. Other coded variables were related to general study characteristics (e.g., country of origin) or relevant to planned moderator analyses (e.g., type of encoding; see Supplementary Material A for all coded measures). Sleep stage specific information (i.e., %SWS or REMS) was only coded if such information was derived from polysomnography (PSG). Additionally, all study authors were contacted to obtain correlations between emotional and neutral memory performance for the sleep and the wake groups separately to increase the precision of the effect size estimation.

## 7. Meta-analysis

### 7.1 Effect size calculation

Given that several studies (23 of 34) did not include a pretest, we used the post-sleep difference between emotional and neutral recognition memory as primary outcome measure. All calculations were based on sample sizes, means, and variances (*SDs/SEMs*) reported for emotional and neutral memory performance. If separate means for positive and negative stimuli were reported, a weighted average was computed to acquire a composite measure of emotional memory, since the number of studies investigating positive stimuli was insufficient to allow for separate analyses. Moreover, if studies reported more than one group that underwent post-encoding sleep and only one group that remained awake, the weighted mean average of both sleep groups was calculated, as these manipulations were not of primary interest for the present meta-analysis. Thus, we were able to include all data on sleep groups without including the same wake group twice.

The effect size calculation was adapted from Morris [49] and required the correlation between neutral and emotional recognition memory performance for the sleep and the wake groups seperately. A meta-analysis on all available correlations was performed to aquire a mean estimate of the correlation between both memory types (*k_sleep_* = 17, *k_wake_* = 11) resulting in mean correlations of *M(r)_sleep_* = 50 [.38 ≤ *M(r*) ≤ .62] and *M(r)_wake_* = 63 [.55 ≤ *M(r*) ≤ .72] (see Supplementary Material B for details). These mean correlations were used to substitute 19 correlations missing in included samples. Afterwards, the effect size per sample was calculated using *R* [50] and the *metafor* package [51] by subtracting the emotional memory benefit for the wake groups from that of the sleep groups. This resulted in a *Standardized Mean Change (SMC*) measure per study that reflected the between-group difference in emotional-memory benefit with estimates > 0 describing an increased emotional memory benefit in the sleep compared to the wake group. Critically, using this approach all sleep versus wake comparisons were treated as between-subject designs. Since only a minority of studies (4 of 22 studies that included wake groups) employed within-subject designs, this was deemed the more appropriate and conservative approach. To increase our data basis, we also included studies that did not contain a wake group (12 of 34 studies). Whenever necessary we estimated the outcome of the respective wake groups by meta-analyzing the emotional memory-benefit in wake groups for whole night and nap designs separately (see Supplementary Material C for details). Thus, 12 wake groups were estimated (3 for nap designs, 9 for whole night designs). All analyses used maximum likelihood estimations and relied on random-effects models that allow for true between-studies variation and for inferences beyond the included samples (i.e., to the wider population) [52]. *SMC* was used as an estimate of the populations effect and its 95% confidence interval (CI) as an indicator of its significance. Residual heterogenity of study effects was assessed by means of *τ*^2^, Cochran’s *Q* statistic, and *I*^2^, which expresses heterogeneity as a percentage ranging from 0 to 100 [53]. All analyses are presented with and without the wake group estimation.

### 7.3 Outliers and influence analyses

Outlier and influence analyses relied on studentized deleted residuals (SDRs), Cook’s distances (CD), and covariance ratios (COVRATIO). SDRs below and above ± 1.96 [54], CD values above .45 [55], and COVRATIOs below 1 [54] were considered outliers.

### 7.4 Moderator analyses

The impact of a-priori determined moderators was tested using subgroup meta-analyses for categorial variables (e.g., nap vs. whole night designs) and meta-regression for continuous (e.g., %REMS) moderators [51]. In the former case, non-overlapping 95% CIs of different moderator levels indicate a significant effect. In meta-regression, the significance of the moderator effect is assessed using the *QM* statistic.

### 7.5 Bias assessment

#### External bias

Results of meta-analyses may overestimate the true population effect due to publication bias [56]. To reduce its potential impact, different search strategies were used to include unpublished data (see 2.1). Moreover, the influence of publication bias was statistically assessed using funnel plots and rank correlations (Kendall’s *τ*) to assess funnel plot symmetry. A significant rank correlation test can be interpreted as indicative of a non-normal distribution of effect sizes around the mean effect, reflective of publication bias. In addition, we used the trim-and-fill method to statistically correct for a potential influence of publication bias [57]. In absence of normally distributed effect sizes, the trim-and-fill algorithm adds “missing” effects and computes a new, unbiased, meta-analysis.

#### Internal bias

Meta-analytical findings may also be biased by insufficient study quality such as flaws in study design, conduction, analysis or reporting [58]. Since standard internal-bias-assessment checklists for intervention studies were not applicable, we developed an 11-item quality checklist based on state-of-the-art criteria in sleep-memory research (e.g., degree of experimental control, assessment of sleepiness; see Supplementary Material D for the checklist). To statistically investigate the impact of study quality on the effect size estimation, studies were divided into two groups using a median split (low vs. high quality) and differences were assessed by means of subgroup-moderator analyses.

## 3. Results

### 3.1 Sample description

The final meta-analysis was based on 22 sleep and wake comparisons based on 26 samples consisting of 1,059 observations (*n_sleep_* = 596, *n_wake_* = 463). Including samples without wake group, the meta-analysis comprised 34 sleep wake comparisons based on 38 samples consisting of 1,382 observations (*n_sleep_* = 919, *n_wake_* = 463). All studies were published between 2007 and 2019. The weighted mean sample age was 22.46 years (*SD* = 3.31), and 22.25 years (*SD* = 3.00) when including samples without wake groups. The mean percentage of female participants was 57.54 (*SD* = 13.84), 57.62%, respectively (*SD* = 16.49). Six samples (including samples without wake groups: 9) underwent a nap manipulation, while 14 samples (23, respectively) underwent nocturnal sleep. One comparison involved a sleep deprivation group [37], while all others used daytime wake conditions. All included studies, their sample characteristics, and moderator-relevant information are displayed in Supplementary Material E.

### 3.2 Tests of basic assumptions: Benefit of emotional versus neutral item memory and sleep versus wakefulness

As a prerequisite for the assessment of potential differential effects of sleep compared to wakefulness on emotional versus neutral memory, two meta-analyses investigated the effect of stimulus valence (emotional vs. neutral) on memory consolidation in the sleep and wake groups separately (for details see Supplementary Material C). Analyses showed a higher memory performance for emotional stimuli post-sleep (*k* = 34), *M(SMC*) = 0.41, [0.14 ≤ *M(SMC*) ≤ 0.69], and post-wake (*k* = 22), *M(SMC*) = 0.59, [0.24 ≤ *M(SMC*) ≤ 0.95], compared to neutral stimuli.

Two further meta-analyses were performed to assess whether sleep compared to wakefulness enhanced memory performance for neutral and emotional stimuli, by comparing post-sleep and post-wake recognition memory for neutral and emotional stimuli, separately (for details see Supplementary Material F). These analyses yielded significant *Standardized Mean Differences (SMD*) reflecting better recognition performance following sleep compared to wakefulness, for neutral (*k* = 22), *M(SMD*) = 0.47, *p* < .001, [0.30 ≤ *M(SMC*) ≤ 0.63], and emotional stimuli (*k* = 22), *M(SMD*) = 0.47, *p* < .001, [0.29 ≤ *M(SMC*) ≤ 0.65].

### 3.3 Differential effect of sleep on emotional memory: Overview of study results

The forest plot displays the effect sizes and CIs of all samples including those without wake groups (Figure 3). The effect sizes of this analysis ranged from −1.11 to 1.16 with ten out of 34 studies (29.41%) reporting a positive effect, with a numerically positive *SMC* equating to better memory for emotional versus neutral items in the sleep versus the wake group (i.e., a selective effect of sleep on emotional memory consolidation).

**Figure 3.**
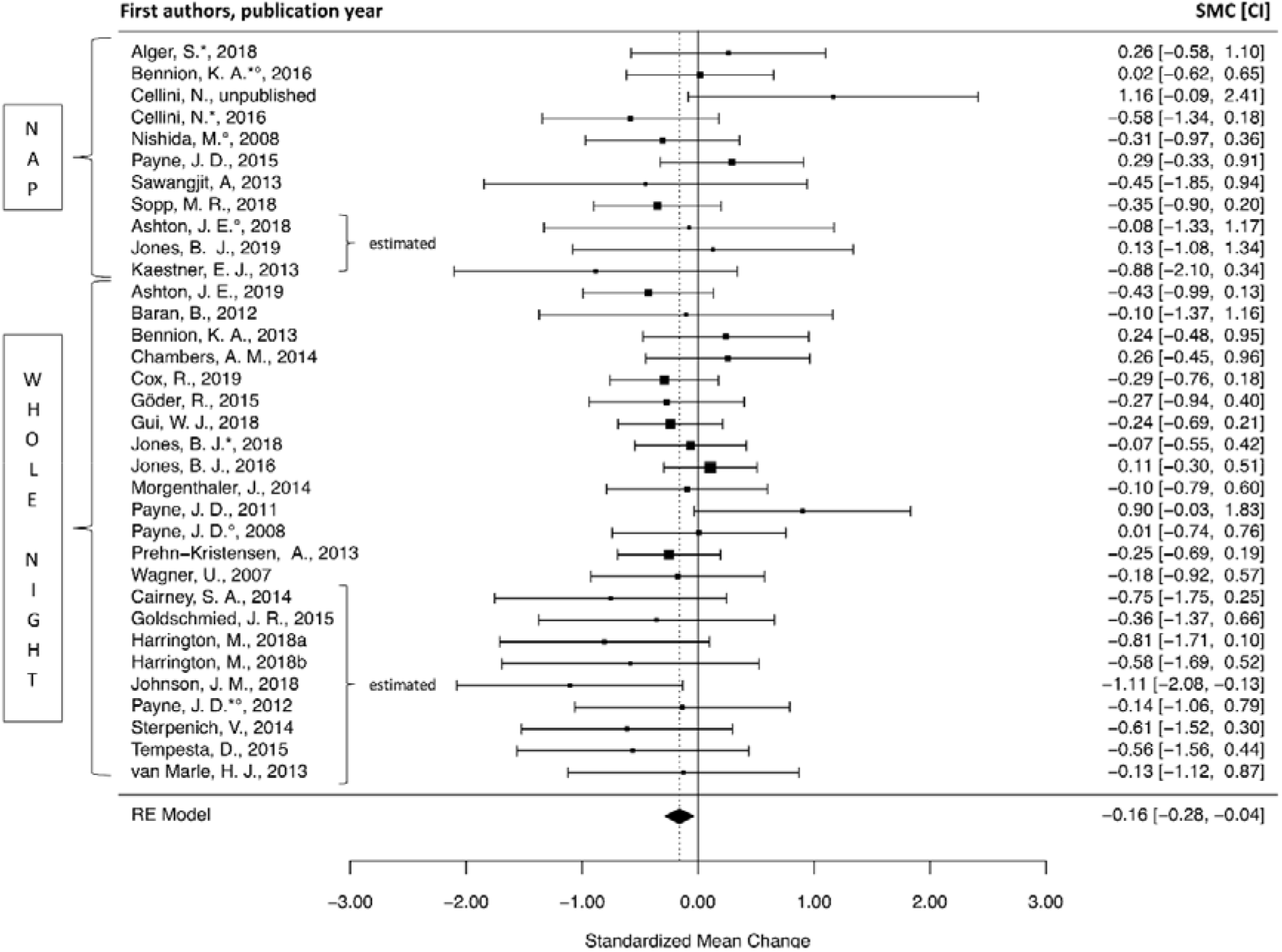
Forest plot displaying Standardized Mean Changes (squares) and confidence intervals (whiskers) of all studies including those without a wake group, as well as the overall effect (diamond). The size of each study’s square represents the precision of the study’s estimate. * marks studies that included more than one sleep group (effect sizes were calculated as weighted mean of the sleep groups vs. the wake groups). ° marks studies where data was extracted using *WebPlotDigitizer*. RE = random effects.

The forest plot showing the effect sizes without the estimation of 12 wake groups revealed a similar pattern of results (see Supplementary Material G). The effect sizes of this analysis ranged from −0.58 to 1.16 with nine out of 22 studies (40.09%) reporting a positive effect in terms of a preferential consolidation of emotional versus neutral memory after sleep as opposed to wakefulness.

#### 3.4 Meta-analytical results

Table 1 presents the results of the main meta-analyses using a random-effects model. The analysis based on all studies that included a wake group yielded a non-significant mean population effect of *M(SMC*) = −0.09, [−0.22 ≤ *M(SMC*) ≤ 0.04]. Including the estimation of 12 wake groups, the same analysis showed a significant mean population effect of *M(SMC*) = −0.16, [−0.28 ≤ *M(SMC*) ≤ −0.04]. Thus, contrary to the widely reported hypothesis, the results based on the wake group estimation revealed a significant benefit of wakefulness as opposed to sleep for emotional versus neutral memory. In both analyses, there was no evidence of heterogeneity of effect sizes, indicated by *τ^2^* = .00, a non-significant *Q* statistic (*p* > .538), as well as *I^2^* = .00. The absence of heterogeneity supports the generalizability of findings beyond the included studies to the population [59].

**Table 1.**
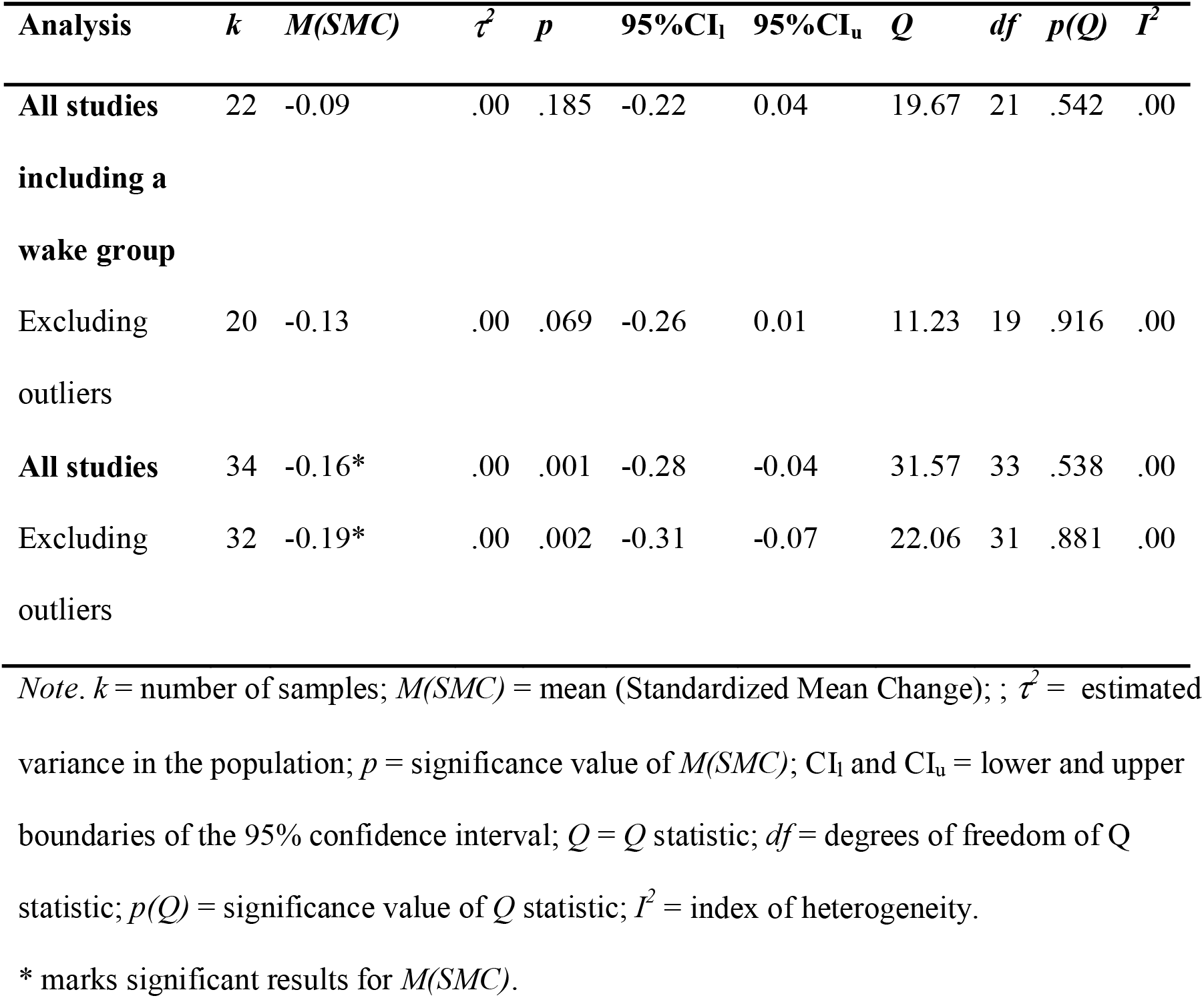
Results of the main meta-analyses

### 3.5 Outlier and influence analyses

An outlier analysis of all studies that included a wake group identified two outlying samples based on SDRs, which were above 1.96 [54] (see Supplementary Material H for outlier statistics). Thus, both main analyses were re-calculated excluding the outliers (see Table 1). However, the results were identical as reflected in strongly overlapping CIs. Considering the general absence of heterogeneity, the outliers were included in subsequent moderator analyses.

### 3.6 Moderator analyses

Given the absence of heterogeneity, it is debatable if moderator analyses should be performed as these aim to explain heterogeneity. However, it is suggested to perform a-priori planned moderator analyses even in presence of homogeneity to explore remaining between-study variance [60].

As shown in Supplementary Material I, planned analyses did not reveal a significant moderating effect of valence, type of sleep manipulation (nap vs. whole night), pre-sleep memory test, type of learning (incidental vs. explicit) or study quality (high vs. low). Finally, no moderating effects were identified for publication year, sample age, gender imbalance (expressed as %female), TST, %REMS or %SWS as indicated by non-significant *QM* statistics (all *p*s > .525) and *R^2^* of .00 (see Supplementary Material I).

### 3.7 Publication bias

Visual inspection and non-significant rank correlations (without wake group estimation: Kendall’s *τ* = .19, *p* = .239; with wake group estimation: Kendall’s *τ* = −.08, *p* = .498) indicated no asymmetry of the funnel plots. Thus, there was no evidence for a substantial influence of publication bias on the meta-analytical results and no need to correct its impact using trim-and-fill analyses.

### 3.8 Additional meta-analysis: REMS versus SWS

We conducted an additional meta-analysis on the split-night designs and REM/SWS deprivation studies identified by the literature search (details on the meta-analysis are provided as Supplementary Material J). The analysis (*k* = 5) comparing early sleep/REMS deprivation and late sleep/SWS deprivation groups revealed a significant benefit of late sleep/SWS deprivation on emotional memory consolidation, *M(SMC*) = 0.35 [0.05 ≤ *M(SMC*) ≤ 0.66] in absence of heterogeneity [*Q*(4) = 1,02, *p* = .906, *I*^2^ = .00]. This result did not change when estimating three additional late sleep/SWS deprivation groups (*k* = 8) [33], *M(SMC*) = .39 [0.13 ≤ *M(SMC*) ≤ 0.66]. Figure 4 displays the forest plot.

**Figure 4.**
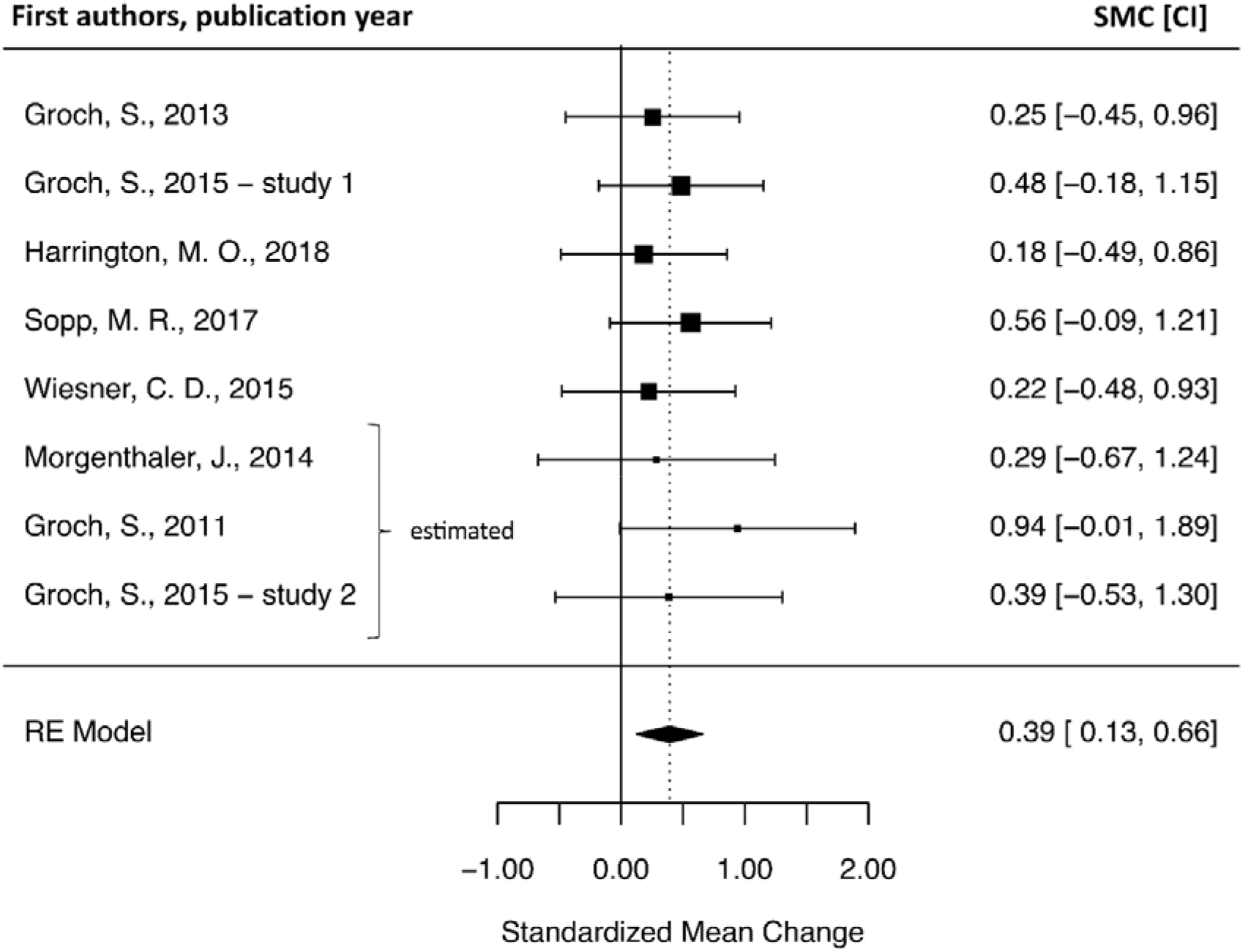
Forest plot displaying Standardized Mean Changes (squares) and confidence intervals (whiskers) of split-night and REM/SWS deprivation studies including the estimation of three late sleep/SWS deprivation groups, as well as the overall effect (diamond). The size of each study’s square represents the precision of the study’s estimate. The forest plot displaying the results without the estimation of one late sleep/SWS deprivation group is provided in Supplementary Material J. RE = random effects.

## 4. Discussion

The present meta-analysis quantified the effects of sleep on emotional memory consolidation based on 34 samples from nap/whole-night studies and 8 samples from split-night/REM/SWS deprivation studies. Tests of basic assumptions confirmed that emotional memory was superior to neutral memory after both wakefulness and sleep. Moreover, both neutral and emotional memory performance were higher after sleep than after wakefulness. However, in contrast to theoretical frameworks, our main analysis did not yield a selective effect of sleep on emotional memory consolidation. Conversely, including a maximum number of studies by estimating wake group effects, we found a larger difference between emotional and neutral memory performance after wakefulness as opposed to sleep, suggesting that processes during wakefulness might be driving the selective retention of emotional memories. Interestingly, our additional meta-analysis of study designs contrasting REMS-rich sleep with SWS-rich sleep yielded a significant effect, indicating a greater difference between emotional and neutral memory performance after REMS-rich sleep than after SWS-rich sleep. Considering our main analysis, this finding suggests that REMS influences emotional memory consolidation without resulting in a greater difference between emotional and neutral memory performance after whole-night sleep as compared to wakefulness.

### 4.1 Effects of sleep versus wakefulness on emotional memory consolidation

Analyzing all studies that included a wake group (*k* = 22) revealed a non-significant negative effect size, indicating comparable differences between emotional and neutral memory performance after sleep and wakefulness. As such, our main analysis confirms the results of a previous meta-analysis [43] while using a more refined analysis strategy (with respect to sample characteristics, outcome measures, and handling of multiple study subsamples) and accounting for the within-subject correlation between emotional and neutral memory performance. This result seems surprising considering that many studies included in our analyses have reported significant sleep-related effects on emotional memory (see e.g., [9,22,24,25]). To ensure comparability of outcome measures, we only included post-sleep/wake memory performance measures in our meta-analysis, preferably collapsed across different response types (i.e., Remember- and Know-responses). However, a significant number of included studies (45.5%) used other indices of memory performance (e.g., including only Remember-responses) in their original analyses, which were calculated in different ways (e.g., % retained or pre-post-difference scores). Hence, the effect sizes used for the current analysis do not necessarily align with those reported in individual studies. Overall, this divergence indicates strong variability of analysis procedures, which may result from the lack of gold-standard procedures in the sleep-memory field. Moreover, the fact that we found a homogeneous effect size estimate (*I^2^* = .00) using equivalized outcome measures could indicate that the use of different outcome measures drives variability of study findings. Beyond differences in analysis procedures, studies varied in multiple design features (e.g., sleep manipulation, stimulus valence, and learning instructions). We aimed to account for these procedural differences by conducting moderator analyses. None of these analyses yielded significant effects, which is unsurprising given the lack of significant heterogeneity of effect sizes (*I^2^* = .00). Hence, our results suggest that variance in outcome measures – rather than procedural differences – may influence the magnitude of reported effect sizes.

### 4.2 Effects of wake retention on emotional and neutral memory performance

To include as many samples as possible, we conducted an extended analysis of *k* = 34 studies by estimating wake group parameters in a separate meta-analytical procedure. This analysis yielded a significant effect size estimate, indicating that differences between emotional and neutral memory performance were larger post-wakefulness than post-sleep. Although caution is warranted in interpreting this finding, it is interesting to note that the direction of the effect size aligns with our conservative analysis (*k* = 22) and the effect sizes reported for negative recognition memory by Lipinska et al. [43]. Hence, this significant effect may reflect an increase in statistical power due to the extended pool of samples. Although rather small in magnitude, this effect could indicate that wakefulness – rather than sleep – promotes superior memory for emotional stimuli. When interpreting this finding, it must be considered that our tests of basic assumptions revealed higher memory performance after sleep in both valence categories as well as higher emotional as compared to neutral memory after both sleep and wakefulness. Taken together, these findings suggest a permissive – rather than active – effect of wakefulness on emotional memory retention. A theoretical framework that would predict such a pattern of results is the emotional binding model [61]. The model assumes that emotional memories are supported by amygdala-dependent bindings whereas neutral memories are supported by hippocampus-dependent bindings. These binding mechanisms are assumed to account for differences in emotional and neutral memory retention across time: Whereas hippocampus-dependent bindings are assumed to be highly susceptible to interference, amygdala-dependent bindings are considered resistant to interference. Consequently, stronger decrements of neutral as opposed to emotional memories are believed to emerge across time. Assuming that sleep may not just actively support consolidation of emotional and neutral memories but also passively protect hippocampus-dependent bindings from retroactive interference (due to the absence of sensory processing during sleep [62]), one would predict a smaller difference between emotional and neutral memory performance after sleep than after wakefulness as found in the current analysis. However, these assumptions should also be reflected in higher neutral memory performance after sleep as compared to wakefulness, which cannot be tested based on our extended sample (*k* = 34). Thus, more research is required to confirm this interference-based account. If confirmed, such an account would accommodate previous findings demonstrating that emotional memory is enhanced after shorter retention intervals without sleep (see e.g., [63]).

### 4.3 Effects of REMS versus SWS on emotional memory consolidation

Moderator analyses did not reveal an impact of relative REMS or SWS duration on effect sizes in our main analysis. However, our additional meta-analysis of studies contrasting REMS-rich and SWS-rich sleep yielded a significant effect size indicating that the difference between emotional and neutral memory performance was higher after REMS-rich sleep as compared to SWS-rich sleep. This finding is consistent with several theoretical frameworks postulating that emotional memories are reactivated during REMS [27–29]. These frameworks assume that selective reactivations of emotional memories are reflected by the propagation of REMS theta oscillations that arise from synchronized activation in the neural structures involved in the encoding of emotional memories (i.e., hippocampus and amygdala). While some studies have provided supportive evidence of this hypothesis [24,26], others have reported inconsistent findings (see e.g., [34]). Although the present analyses cannot address the underlying mechanisms of emotional memory consolidation, our findings indicate an enhancing effect of post-encoding REMS on emotional memory performance. However, our main analyses do not indicate that this enhancement is evident after a whole night of sleep. Based on these results, it could be speculated that REMS enhances emotional memory consolidation whereas SWS enhances neutral memory performance resulting in similar performance levels after a whole night of sleep [64]. Future research should investigate this notion. Moreover, given that we could only include a limited number of studies in this additional analysis (*k* = 8), future meta-analyses need to confirm our findings using more samples. However, despite the limited number of samples and the fact that we collapsed REM/SWS deprivation and split-night designs in one analysis, our conservative and extended analyses both revealed a homogenous effect size estimate (*I^2^* = .00), supporting the reliability of our findings. Nevertheless, the neurophysiological basis of this effect and its direct association with REMS require further study. As most of our samples came from split-night-study designs, it is possible that higher emotional as compared to neutral memory performance may have resulted from circadian effects (but see [65]). That is, the increase of cortisol levels – rather than REMS – during late night sleep might have influenced consolidation [66].

### 4.4 Limitations and implications for future research

Several limitations of the current analyses need to be considered. As it was necessary to equivalize effects for meta-analytical effect size estimation, we chose our outcome measure (post-sleep/wake memory performance) based on the data that was available for most studies. As a result, we were not able to examine whether sleep affects pre-post changes in memory performance and memory performance for specific response types (e.g., recollection-based responses). However, the number of studies reporting such response-specific analyses would not have been sufficient to conduct separate analyses. Thus, future meta-analyses need to address this issue based on a larger number of samples. In addition, we focused our analyses exclusively on recognition memory. This decision was based on memory models assuming that recognition memory tests are supported by other retrieval processes than recall tests or associative memory tests [10]. Given the small number of studies and the high variability of outcome measures, we decided against conducting a separate analysis of recall and associative memory tests. However, such analyses should be reconsidered once the pool of empirical studies is sufficient (see also [43]). Moreover, future meta-analyses should examine sleep’s effects on emotional memory in other memory domains. In this regard, several studies indicate that sleep may affect the retention of conditioned fear responses (for reviews see [67,68]). This area of research appears particularly promising since most studies investigating the underlying mechanisms of emotional memory consolidation during sleep have used fear conditioning procedures [31,32]. Overall, the present findings as well as the necessity to equivalize outcome measures to enable a joint meta-analysis illustrate the need for a greater alignment of methodological approaches in the sleep-memory field. Future research should thus define gold standards for study designs and data analysis as it has been previously done in other fields of research (see e.g., [69]).

### 4.5 Conclusions

Our main analyses of sleep/wake effects and our additional analysis of REMS/SWS effects seem to yield diverging findings: On the one hand, we found indications that post-encoding wakefulness results in enhanced emotional as opposed to neutral memory performance. On the other hand, our additional analyses suggest that post-encoding REMS may strengthen emotional memory performance to a greater extent than post-encoding SWS. This result pattern could reflect the complexity of modulatory effects on emotional memory retention: Some of these effects may result in behavioral changes – as evident in enhanced retention of emotional memories across wakefulness – whereas sleep-related effects may be subtler and only emerge if REMS effects are contrasted against SWS effects (i.e., blocking out inference effects). This notion should be explored by future research. Nevertheless, with respect to behavioral outcomes of post-encoding wakefulness and sleep, our results do not support the hypothesis that sleep enhances emotional memory beyond the enhancement observed for neutral memory.

#### Research Agenda

1. Strong variability in data-analysis strategies indicates the need for gold-standard procedures in the sleep-memory field.
2. Future studies should investigate the effects of individual sleep stages (esp. REMS) on emotional memory using well-controlled designs.
3. Future meta-analyses should examine the effects of sleep in other emotional memory domains (e.g., fear conditioning).

#### Practice Points

1. Sleep benefits emotional and neutral memory performance.
2. When contrasted against wakefulness, sleep does not increase the difference between emotional and neutral memory performance.
3. Post-encoding REMS may be associated with a larger difference between emotional and neutral memory than post-encoding SWS.

## Abbreviations

CI: confidence interval
CD: Cook’s distances
COVRATIO: covariance ratios
NREMS: non-rapid-eye-movement sleep
PSG: polysomnography
REM: rapid-eye-movement sleep
SDRs: studentized
SDRs: studentized deleted residuals
SMC: standardized mean change
SMD: standardized mean difference
SWS: slow-wave sleep

## Footnotes

1 Samples were included if sleep deprivation was followed by a recovery night and the overall retention interval did not exceed 48 hours (see e.g., [37]). For studies that did not include a recovery night, we estimated wake group effects (see e.g., [45])

